# From Wandering Thoughts to Mental Stillness: A Minimal Computational Statistical-Mechanical Model of Driven Relaxation

**DOI:** 10.64898/2026.10.11.758288

**Authors:** Soumyajit Karmakar

**Affiliations:** Department of Chemistry, Indian Institute of Technology Kharagpur

**Keywords:** spontaneous thought, meditation, stochastic dynamics, bistability, metastability, Langevin equation, driven relaxation, transition entropy, neural dynamics

## Abstract

Reduced spontaneous thought during meditation is often described phenomenologically, but a mechanistic account should specify how a dynamical system can enter and persist in such a regime. Here we formulate a minimal statistical-mechanical model in which a dimensionless coordinate *q* represents a coarse-grained propensity for spontaneous-thought activity and a slowly relaxing variable *r*(*t*) represents the history-dependent effect of an attentional manipulation. The model combines a tilted double-well landscape with Langevin noise, allowing a high-*q* and a low-*q* dynamical regime to coexist. We numerically integrate the model under a controlled focusing protocol followed by removal of the drive and quantify low-state occupancy, first-passage times, dwell times, occupancy entropy, and transition entropy. For the chosen dimensionless parameterization, increasing the drive amplitude produces a sharp crossover in low-state occupancy near the deterministic spinodal threshold *A*_sp_ ≃0.0148. At noise intensity *D* = 0.01, the probability of occupying the low-*q* regime at drive removal increases from 0.094 at *A* = 0 to 0.994 at *A* = 0.02, while the mean first-passage time decreases from 48.0 to 13.5 model time units. Noise produces a non-monotonic effect: for *A* = 0.01, low-state occupancy at drive removal is only 0.033 at *D* = 0.0025, rises to 0.842 at *D* = 0.02, and falls to 0.718 at *D* = 0.04. Strong driving also reduces the effective number of occupied states while changing transition entropy much less, illustrating why a single entropy measure cannot identify thoughtlessness. These results do not establish that human neural dynamics obey this model. Instead, they show that a driven stochastic bistable system can generate the qualitative sequence of focusing, switching, relaxation, and metastable low-activity residence, and they provide quantitative predictions that can be tested against time-resolved neural recordings and independent reports of subjective experience.

## Introduction

A thought rarely exists in isolation. During ordinary waking life, one thought may evoke a memory, an imagined future, an emotion, or another unrelated thought [1]. The mind can therefore be viewed as a system that continually moves between different cognitive states [2]. If *A, B, C*, … denote coarse-grained cognitive states, spontaneous thought can be represented schematically as

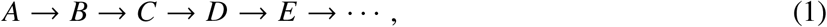

where the sequence need not follow a fixed order and transitions may depend on memory, emotion, sensory input, and other internal processes.

Meditation may alter this dynamical organization. Attention can repeatedly return to a selected object, such as the breath, producing a more restricted pattern of transitions:

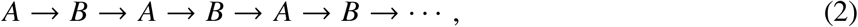

where *A* represents the intended attentional object and *B* a transient deviation. This schematic sequence does not imply that meditation consists of only two states, nor does returning attention to the breath imply complete thoughtlessness. Nevertheless, it raises a fundamental question: *can repeated restriction of cognitive transitions eventually lead to a persistent regime in which spontaneous thought becomes substantially less frequent while wakeful awareness is maintained?* [3]

This question is relevant to several established research areas. Spontaneous thought research emphasizes both the contents of thoughts and the transitions between them [1], while meditation studies have reported changes in large-scale brain networks, including the default mode network [2, 3]. Wakeful thoughtless emptiness has also been investigated electrophysiologically, with reported differences from focused attention, open monitoring, and resting conditions [4]. Neural attractor models provide a framework for describing persistent dynamical states, transitions between regimes, and responses to perturbations [5, 6]. Related computational and theoretical studies have examined meditation in relation to whole-brain dynamics, mind wandering, attention, self-reference, nondual awareness, and consciousness [7–25]. However, studies of meditation and neural complexity report heterogeneous changes in entropy, variability, and dynamical organization, cautioning against assuming that meditation universally corresponds to reduced neural entropy or complexity [26, 27].

The central problem is therefore not the lack of evidence for meditation-related changes or the absence of research on thoughtless experience. Rather, existing approaches do not, by themselves, provide a minimal quantitative model in which attentional history, stochastic fluctuations, switching, and post-manipulation persistence can be varied and tested systematically. In particular, the sequence

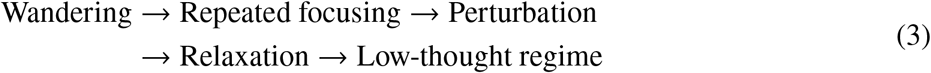

has not been established as a unified, experimentally tested mechanism.

The present work addresses this gap by constructing and simulating a minimal stochastic dynamical model in which these quantities are explicit model observables. The question is therefore not whether meditation affects brain dynamics, but whether a specific dynamical transition can provide a useful and falsifiable model for the transient reduction of spontaneous thought. As a theoretical chemist, I use mathematical models to connect microscopic dynamics with observable behaviour. The equations presented here are therefore modelling tools, not claims that the brain obeys the same physical laws as a chemical reaction or a system at thermodynamic equilibrium. Each assumption is stated explicitly to distinguish what follows mathematically from what remains to be tested experimentally. Within this framework, we test whether a low-spontaneous-thought regime can emerge as a stationary or metastable state of the model, rather than assuming that it must do so. A central aim of the analysis is to distinguish such neural stationarity from the subjective experience of thoughtlessness.

## Model and Methods

### A minimal coarse-grained dynamical model

The objective is to convert the qualitative driven–relaxation hypothesis into a model that can be simulated and falsified. We deliberately use the smallest model that contains the ingredients required by the hypothesis: (i) two candidate dynamical regimes, (ii) a controllable attentional drive, (iii) a finite response timescale, and (iv) stochastic fluctuations. The variables are phenomenological. In particular, *q* is not identified with a measured brain signal and *r* is not identified with a specific neurotransmitter, circuit, or physical force. The latent coordinate *q* (*t*) represents a coarse-grained propensity for spontaneous-thought activity. Smaller *q* denotes the candidate low-spontaneous-thought regime and larger *q* the candidate high-spontaneous-thought regime. The coordinate evolves according to the overdamped Langevin equation

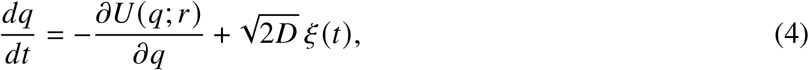

where *D* is the dimensionless noise intensity and ξ (*t*) is Gaussian white noise with

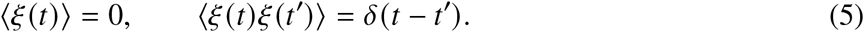

The effective landscape is

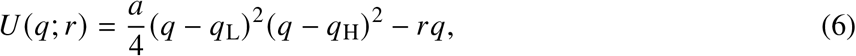

with *q*_L_ < *q*_H_ and *a* > 0. At *r* = 0, the landscape has minima at *q*_L_ and *q*_H_, separated by a barrier at *q*_B_ = (*q*_L_ +*q*_H_) /2. The term −*r q* tilts the landscape without changing the basic one-dimensional character of the model. Negative *r* favours the low-*q* regime; positive *r* favours the high-*q* regime.

The attentional history is represented by a slowly relaxing response variable,

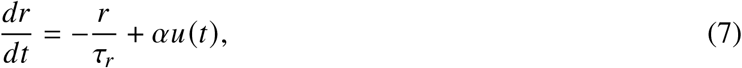

where *u* (*t*) is the imposed attentional drive, s*τ*_*r*_ is its relaxation timescale, and *α* is the drive-response coupling. During the focusing stage we use

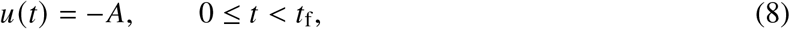

and after the manipulation is removed,

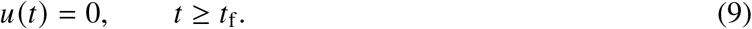

The sign convention is chosen so that focusing favours lower *q*. This is an explicit modelling assumption rather than an empirical statement about the sign of any particular neural variable.

For a constant drive, Eq. (7) gives

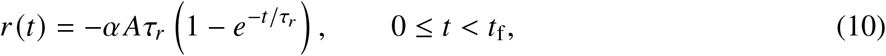

assuming *r* (0) = 0. After drive removal,

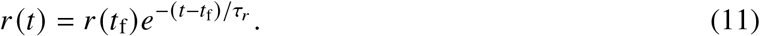

Thus the model contains a finite history-dependent response even after the attentional manipulation has ended. Importantly, this is dynamical persistence, not mechanical momentum.

### Deterministic bistability and the spinodal threshold

The model has a useful analytical control parameter. Define

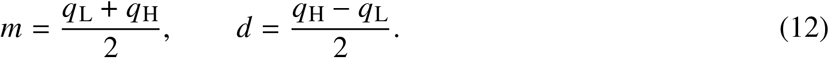

With *x* = *q* − *m*, the stationary points of the deterministic landscape satisfy

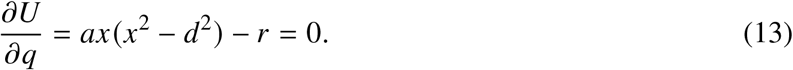

The boundaries of the bistable region occur when a minimum and the intervening barrier merge, requiring both ∂*U*/∂*q* = 0 and ∂^2^*U*/∂*q*^2^ = 0. Since

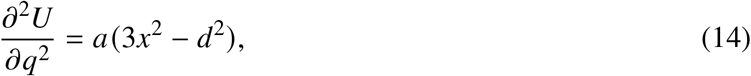

the spinodal positions are

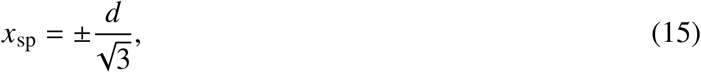

and the magnitude of the critical tilt is

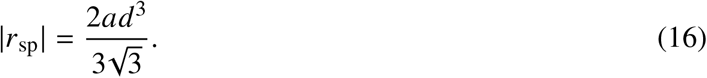

For the parameterization used here,

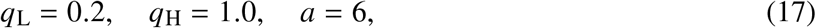

giving

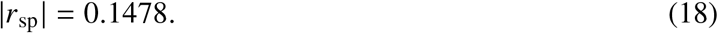

Because the steady response under a constant focusing drive is *r*_∞_ = −*αAτ*_*r*_, the corresponding steady-drive threshold is

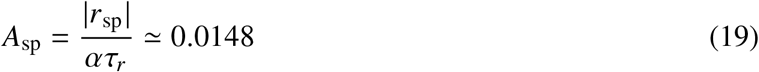

for *α* = 1 and *τ*_*r*_ = 10. This threshold is a property of the model and has no direct empirical interpretation. It nevertheless provides a useful prediction: drive amplitudes near this value should produce a rapid change in the probability of reaching the low-*q* basin.

### Simulation protocol

All simulations use dimensionless model units. The parameter set is summarized in Table 1. The initial condition was *q* (0) = 0.95, placing the system in the high-*q* basin, and *r* (0) = 0. A focusing drive was applied for *t*_f_ = 30 time units and then removed. Each stochastic trajectory was integrated using the Euler–Maruyama scheme,

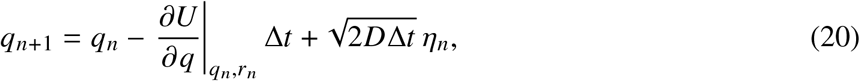

where η_*n*_ are independent standard normal variates. The response variable was propagated using the corresponding Euler step for Eq. (7). A timestep of Δ*t* = 0.005 was used, and trajectories were followed to *t* = 100.

**Table 1:** Parameters of the dimensionless stochastic model.

| Parameter | Value |
| --- | --- |
| $q_L$ | 0.2 |
| $q_H$ | 1.0 |
| $a$ | 6 |
| $\tau_r$ | 10 |
| $\alpha$ | 1 |
| $t_f$ | 30 |
| Total simulation time | 100 |
| $\Delta t$ | 0.005 |
| Initial $q$ | 0.95 |
| Initial $r$ | 0 |
| Low-state threshold | $q < 0.4$ |

For the drive-amplitude analysis, *D* = 0.01 and

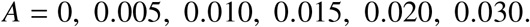

For each amplitude, five independent ensembles of 1000 trajectories were generated. For the noise analysis, *A* = 0.01 and

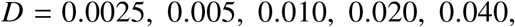

again using five independent ensembles of 1000 trajectories. Reported uncertainties are the standard deviation among these five independent ensemble realizations.

### Observables

The primary observable is the low-state probability

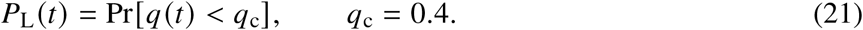

The threshold *q*_c_ = 0.4 is halfway between the low minimum *q*_L_ = 0.2 and the zero-tilt barrier *q*_B_ = 0.6. It is therefore a basin-based classification rather than an arbitrary threshold placed at the midpoint of the two minima.

The first-passage time is defined as

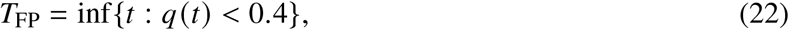

for trajectories initially prepared in the high-*q* basin. The mean of *T*_FP_ is used to quantify how rapidly the focusing protocol transfers trajectories into the low-*q* regime. Low-state dwell time after drive removal is measured from an entry into *q* < 0.4 to the first subsequent exit to *q* ≥0.4.

To connect the stochastic model with the information-theoretic quantities introduced in the original conceptual framework, *q* was also discretized into 20 bins over 0 ≤ *q* ≤ 1.2. The occupancy entropy was calculated as

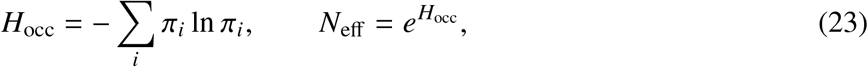

and the transition entropy as

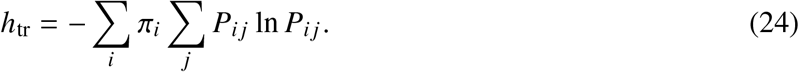

These quantities were calculated from the simulated trajectories between *t* = 30 and *t* = 80, using samples separated by 0.1 time units. Their numerical values depend on the chosen discretization and sampling interval and are therefore treated as model diagnostics rather than universal measures of cognition.

### Numerical reproducibility

The calculations were performed with Python using NumPy, SciPy, pandas, and Matplotlib [28]. The complete simulation protocol, parameter sweeps, random seeds, and tabulated results are supplied with the manuscript files. The stochastic calculations use independent pseudorandom streams for each replicate ensemble. Because the model is dimensionless, the numerical results should be interpreted as relationships among model parameters rather than predictions in seconds, hertz, or other physiological units.

## Results

### A focusing drive reshapes the dynamical landscape

The first result is structural rather than biological: the proposed mechanism can be realized in a minimal stochastic system without assuming a permanent “thoughtless” equilibrium. Figure 1 shows how the focusing response *r* progressively tilts the effective landscape toward the low-*q* basin. At *r* = 0, the two regimes coexist. As *r* becomes negative, the low-*q* minimum becomes deeper and the barrier separating the two regimes becomes smaller. When |*r*| reaches the spinodal value in Eq. (16), the high-*q* minimum disappears in the deterministic model.

**Figure 1:**
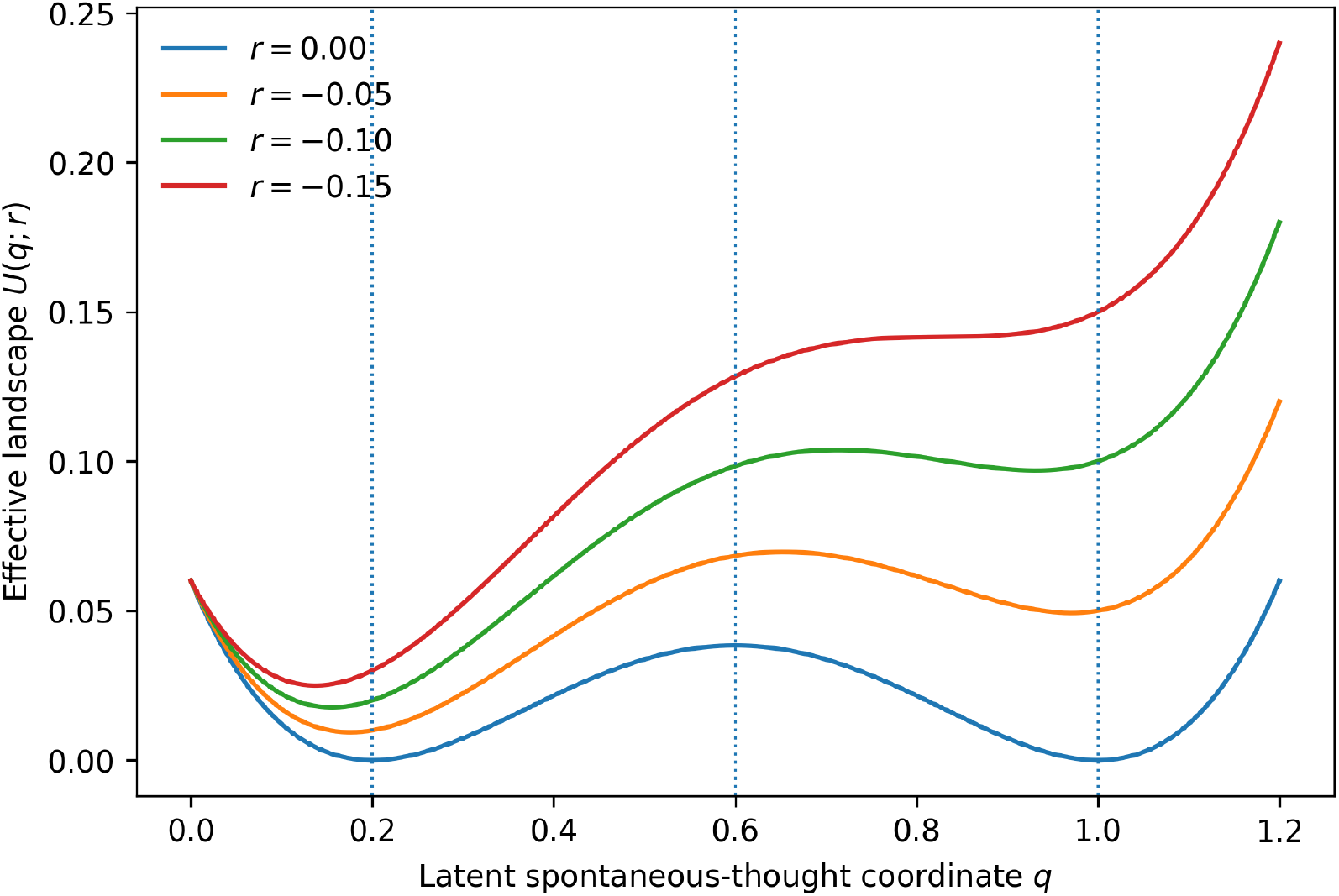
Effective dynamical landscape used in the model. Increasingly negative *r* progressively favours the low-*q* regime. At sufficiently large negative tilt, the high-*q* minimum disappears through a deterministic spinodal transition. The landscape is a phenomenological potential and is not interpreted as a physical energy surface of the brain.

This distinction between barrier crossing and basin disappearance is important. Below the spinodal threshold, stochastic fluctuations are required to move a trajectory between basins. Above the threshold, the high-*q* state is no longer a stable deterministic solution during the driven stage. Thus, the model contains two qualitatively different routes into the low-*q* regime: noise-assisted switching within a bistable landscape and deterministic loss of the high-*q* attractor under sufficiently strong drive.

### Drive removal produces delayed relaxation rather than instantaneous switching

Figure 2 illustrates the history-dependent variable *r* (*t*) for the representative protocol *A* = 0.01, *D* = 0.01. During focusing, *r* approaches a negative value, thereby favouring the low-*q* basin. When the drive is removed at *t* = 30, *r* does not return immediately to zero; instead, it decays exponentially according to Eq. (11). The model therefore produces a finite relaxation interval after the attentional manipulation has ended.

**Figure 2:**
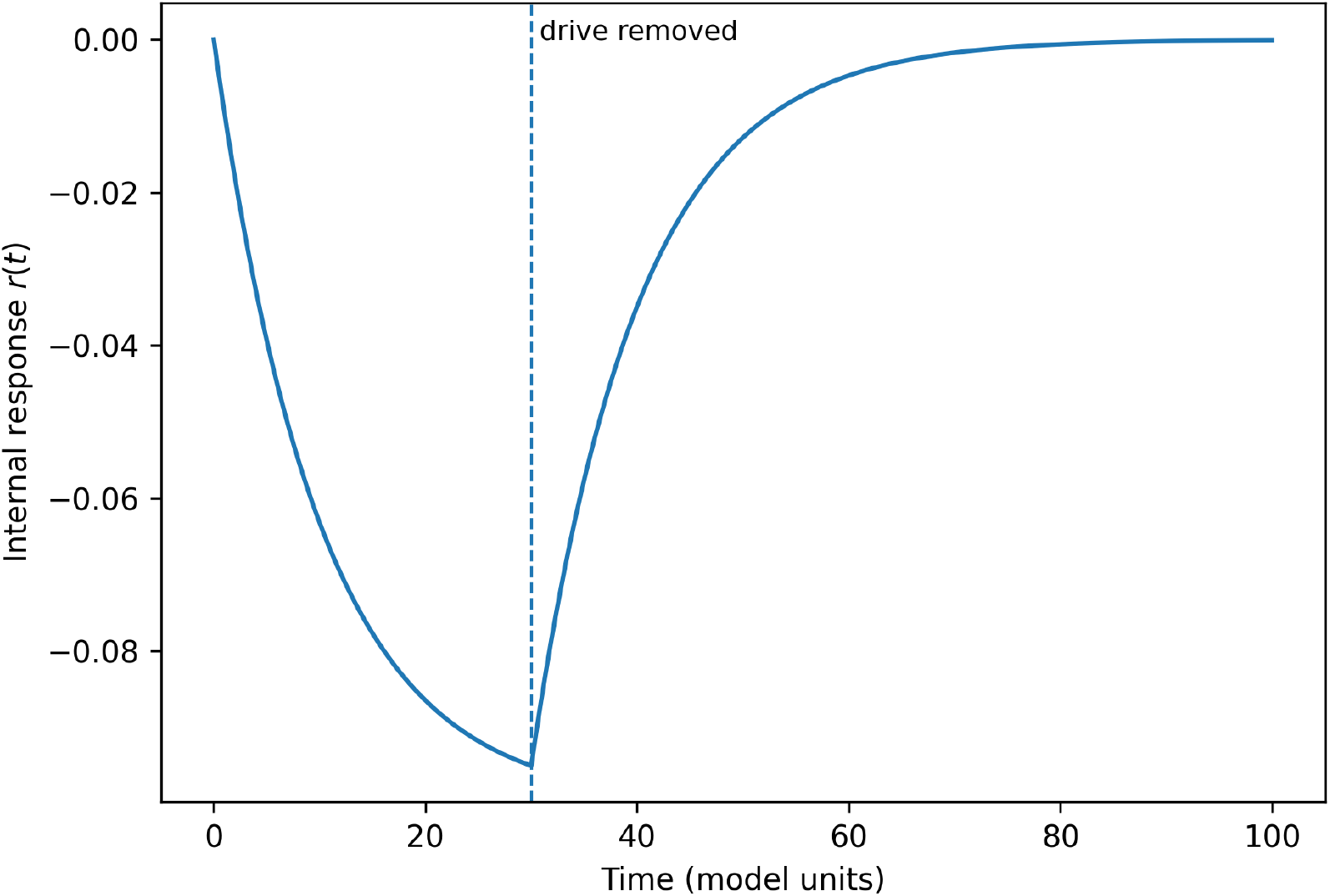
Relaxation of the internal response variable *r* (*t*) after removal of a focusing drive with *A* = 0.01. The vertical dashed line marks drive removal at *t* = 30. The finite decay time is the model’s representation of history-dependent dynamical persistence.

The important point is that persistence of *r* does not itself establish persistence of thoughtlessness. It only creates a time-dependent landscape in which the probability of occupying the low-*q* regime can remain elevated after the external manipulation has ended. The actual state of *q* depends on both the landscape and stochastic fluctuations.

### Drive amplitude creates a threshold-like increase in low-state occupancy

The central computational result is the dependence of low-state occupancy on the strength of the focusing drive. Figure 3 shows *P*_L_ (*t*) for the full amplitude series. In the absence of a drive, spontaneous barrier crossing occurs but the low-*q* regime is occupied only intermittently. Increasing *A* progressively increases low-state occupancy. Near the analytically predicted spinodal amplitude *A*_sp_ ≃0.0148, the response becomes much sharper.

**Figure 3:**
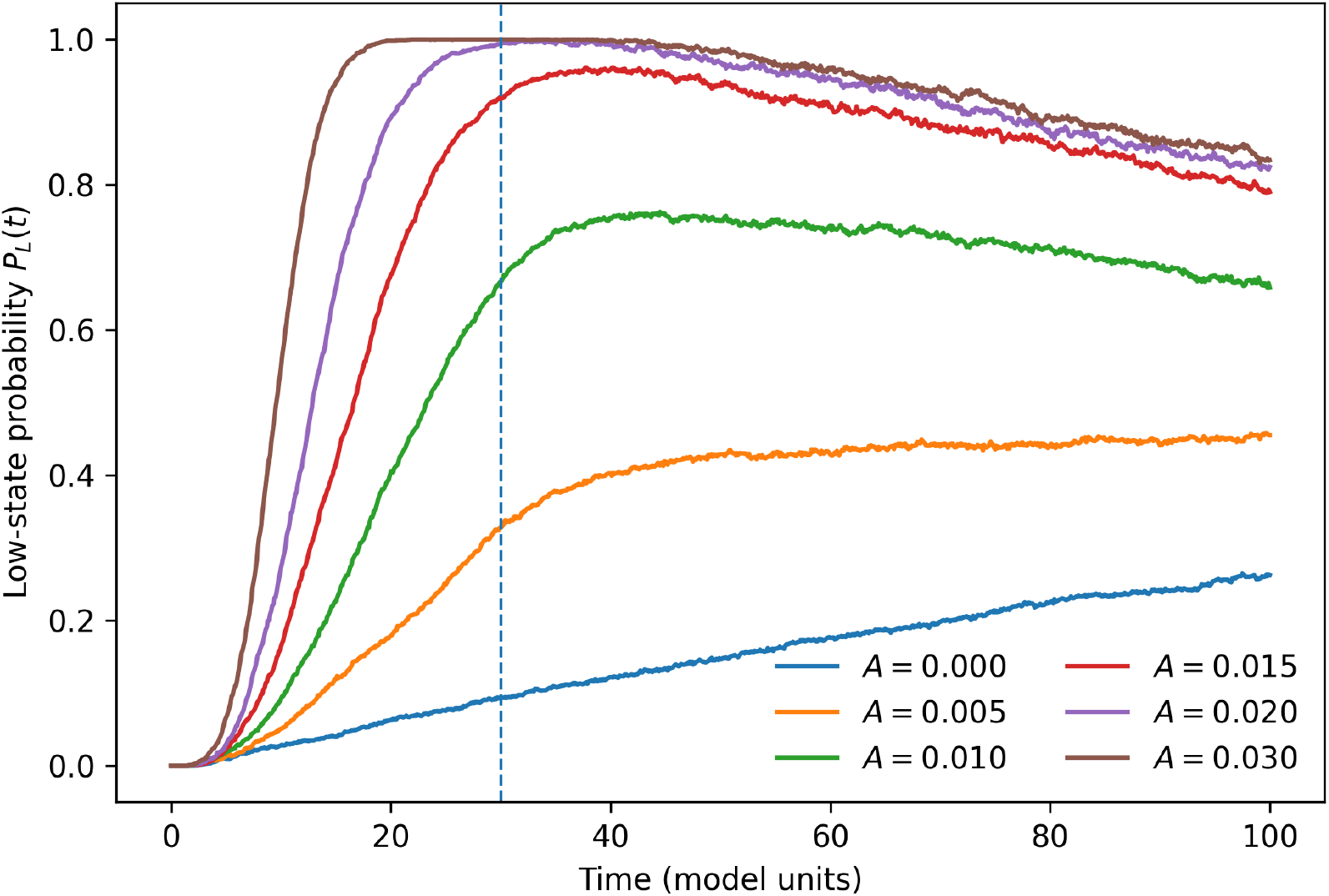
Low-state probability *P*_L_ (*t*) for different focusing amplitudes at *D* = 0.01. The drive is removed at *t* = 30. Stronger focusing produces a larger and more persistent low-*q* population, with a pronounced crossover near the deterministic spinodal amplitude.

**Figure 4:**
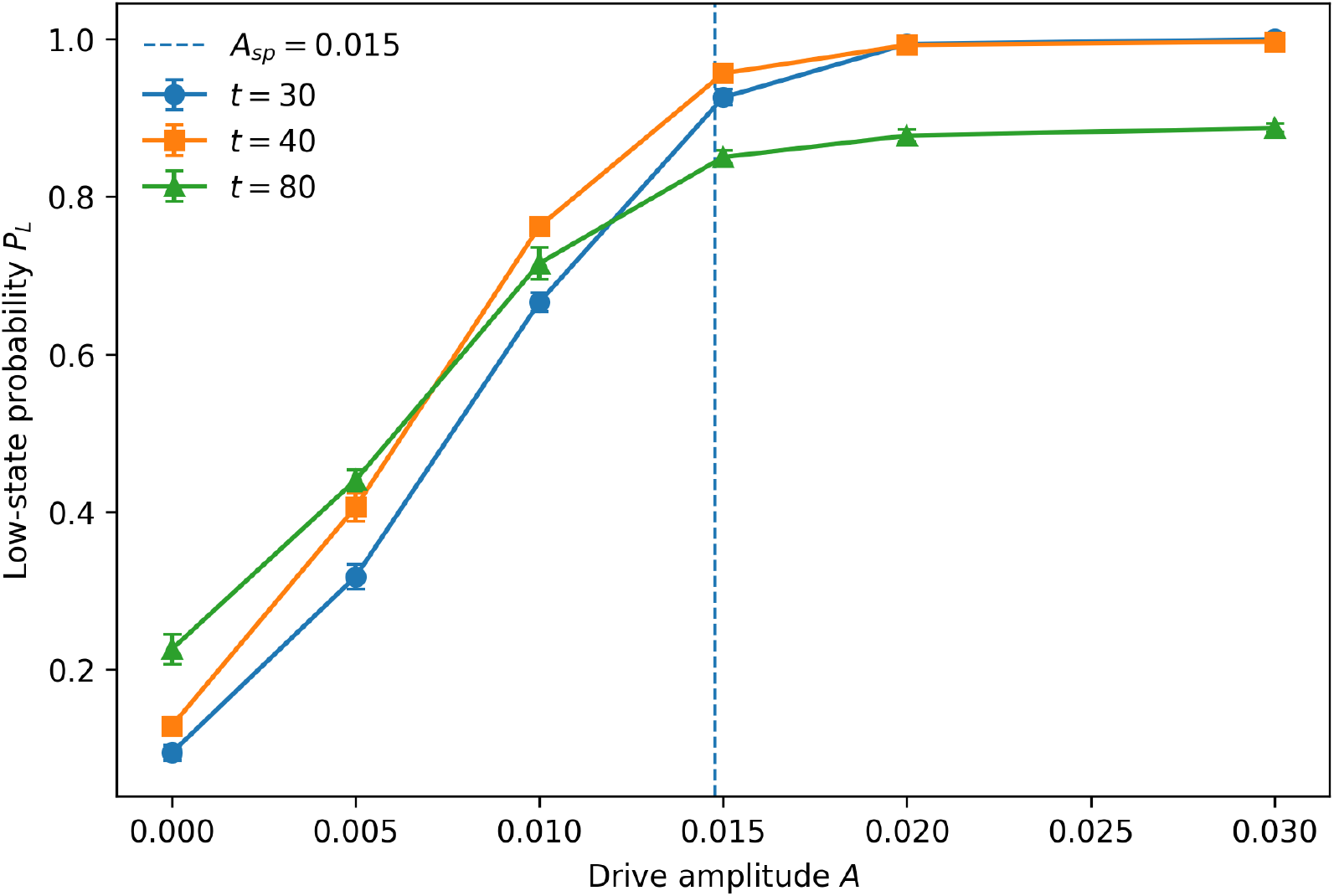
Drive-amplitude dependence of low-state occupancy. Points show means across five independent ensembles and error bars show the standard deviation between ensembles. The vertical dashed line marks the analytically predicted deterministic spinodal amplitude *A*_sp_ ≃ 0.0148.

**Figure 5:**
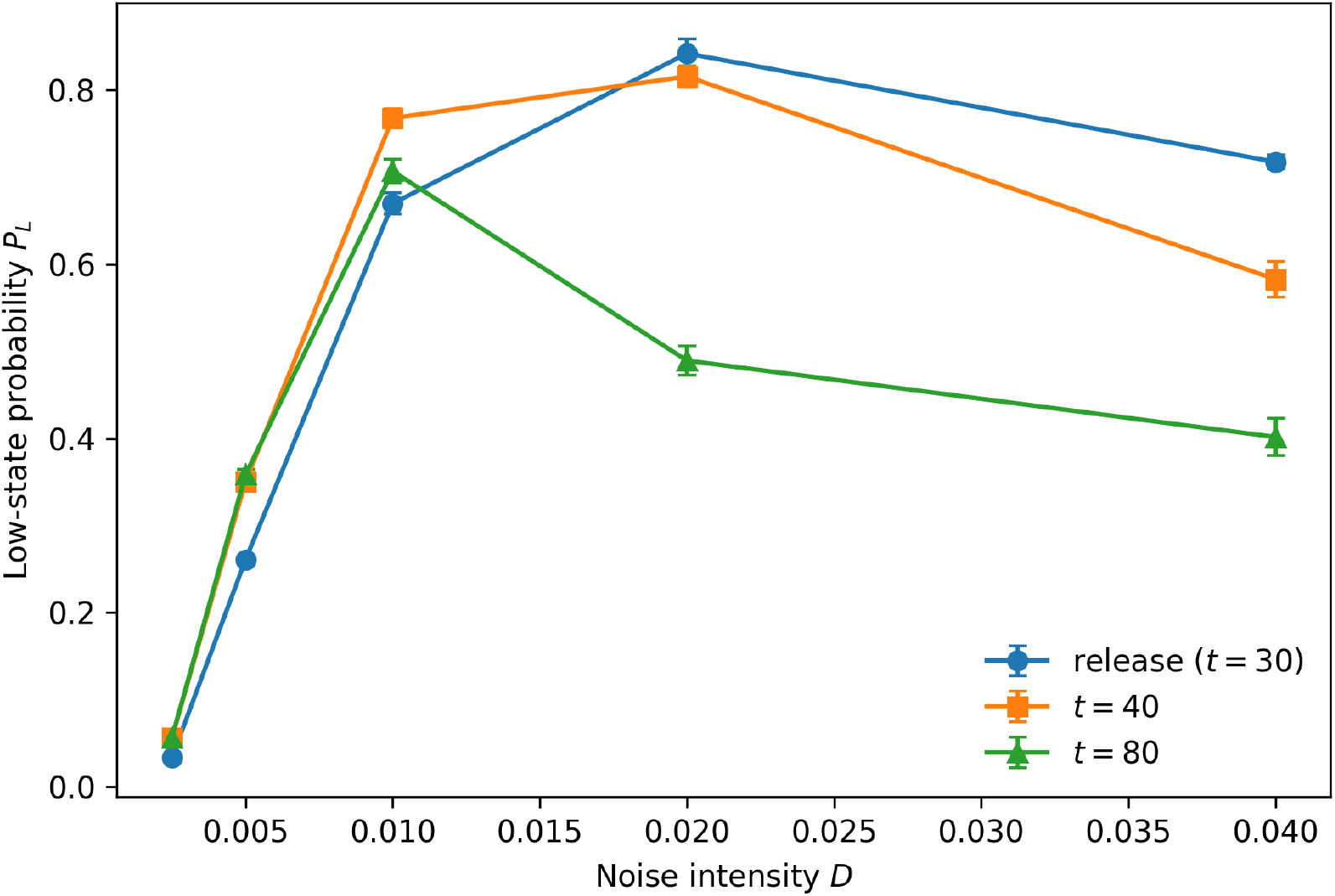
Noise dependence of low-state occupancy for *A* = 0.01. Intermediate noise maximizes access to the low-*q* regime, whereas weak noise suppresses barrier crossing and strong noise reduces persistence. The non-monotonic response is an intrinsic prediction of the stochastic bistable model.

**Figure 6:**
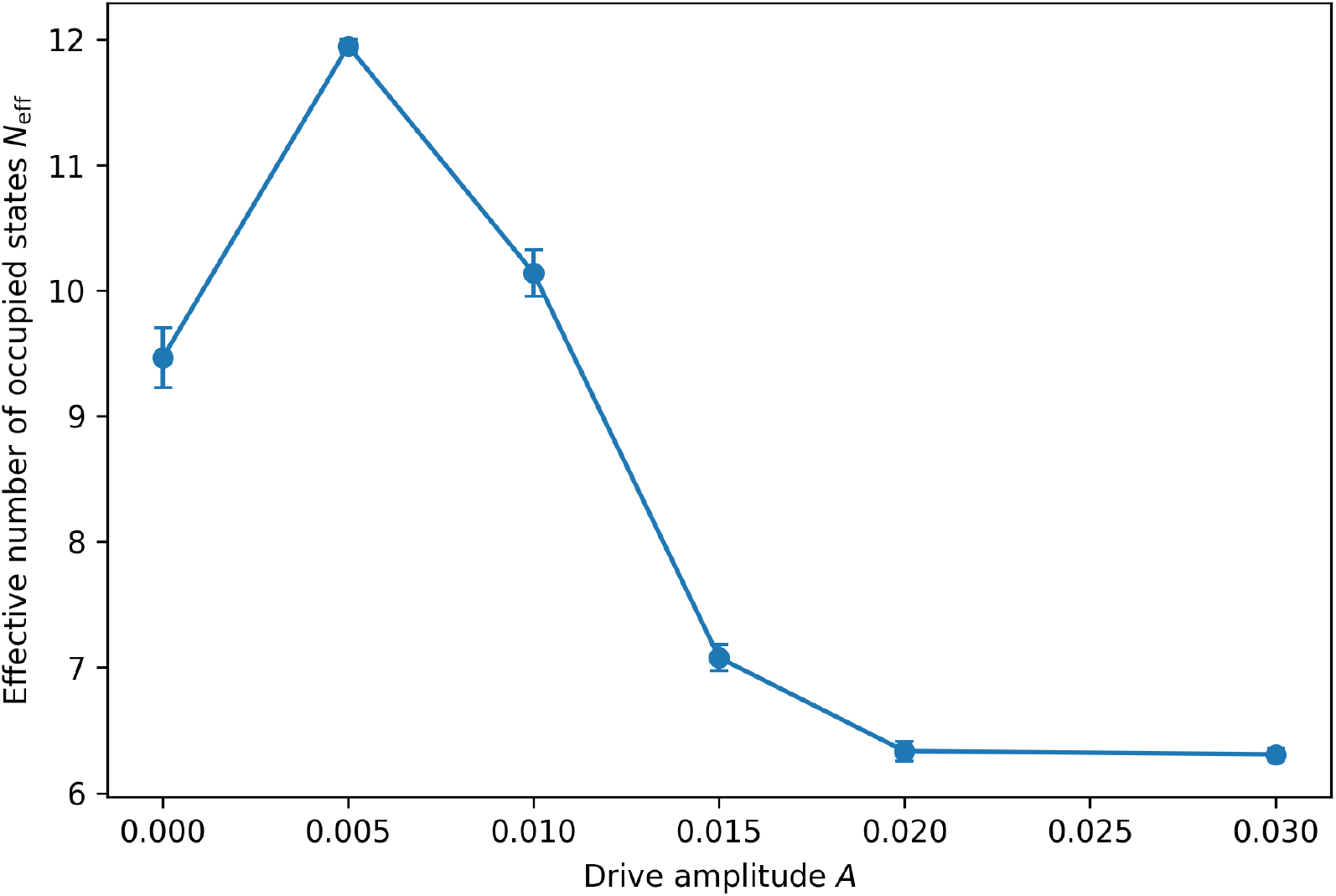
Effective number of occupied coarse-grained states *N*_eff_ as a function of focusing amplitude. Strong focusing compresses the occupancy distribution after the drive has been removed.

**Figure 7:**
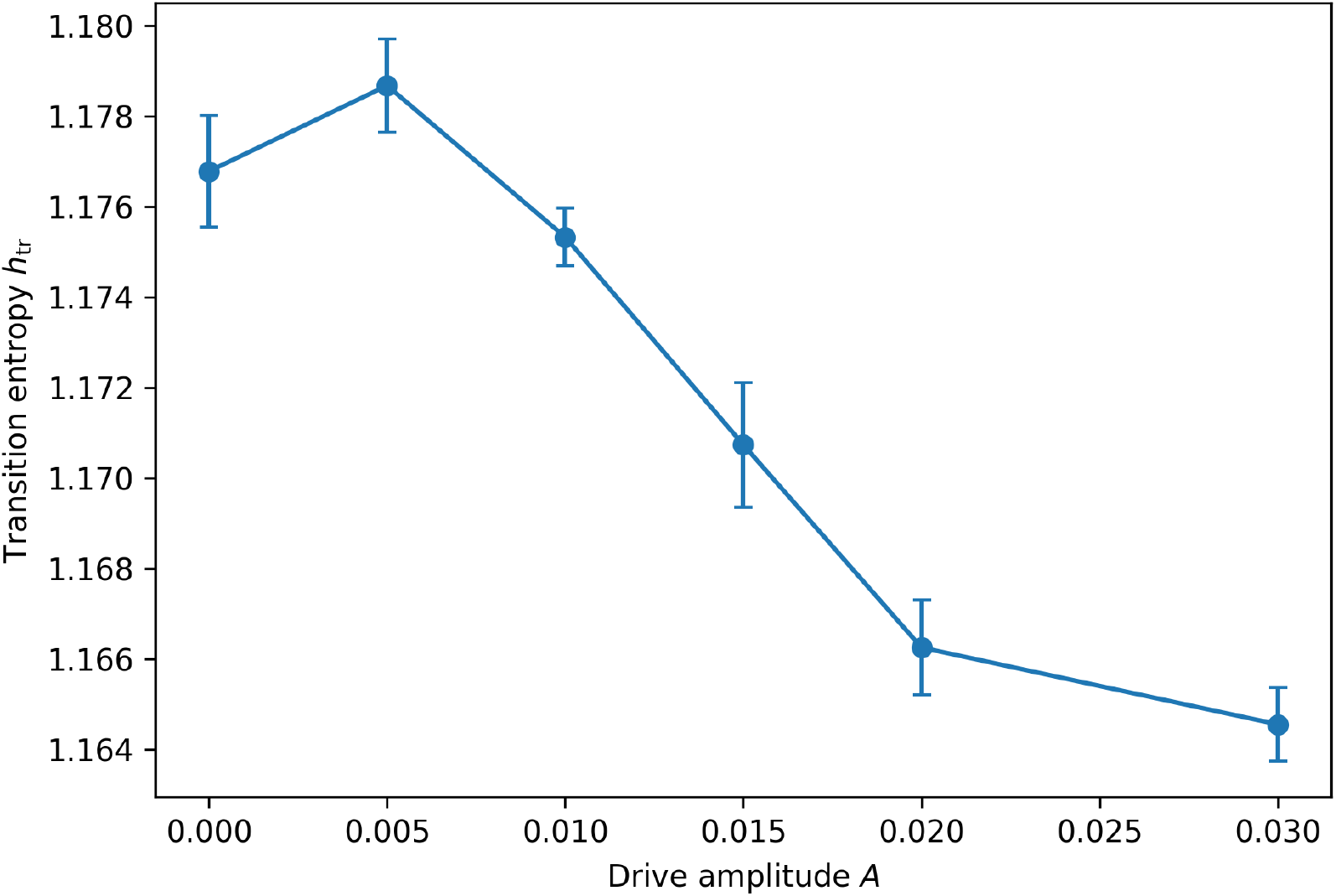
Transition entropy *h*_tr_ as a function of focusing amplitude. The comparatively small change illustrates that reduced occupancy and reduced next-state uncertainty are not equivalent properties.

At *D* = 0.01, the ensemble-averaged low-state probability at drive removal increases from 0.094 ± 0.008 for *A* = 0 to 0.317± 0.010, 0.666± 0.009, 0.926 ±0.007, 0.994± 0.002, and 1.000± 0.000 for *A* = 0.005, 0.010, 0.015, 0.020, and 0.030, respectively. The same ordering persists after drive removal: at *t* = 80, *P*_L_ is 0.225, 0.440, 0.715, 0.850, 0.878, and 0.887, respectively. Thus, the model does not simply produce a transient response locked to the external drive; stronger focusing prepares trajectories that remain preferentially in the low-*q* basin after the manipulation has ended.

The first-passage analysis provides a complementary measure. The mean first-passage time from *q* = 0.95 to *q* < 0.4 decreases from 48.0 model time units at *A* = 0 to 32.5, 23.0, 16.9, 13.5, and 9.5 at the six increasing amplitudes. The decrease is continuous even though the low-state probability changes most rapidly near the spinodal region. This distinction indicates that the model’s threshold-like occupancy change is not merely an artifact of assigning a binary label to *q*; the underlying kinetics also accelerate.

The dwell-time analysis gives the complementary persistence measure. For the same amplitude series, the mean completed low-state dwell time increases from approximately 4.0 model time units at *A* = 0 to 27.2 at *A* = 0.03. Because the model contains noise, the low-*q* state remains metastable rather than permanent. The result therefore supports the narrower statement that a sufficiently strong focusing history can increase both the probability of entering the low-*q* basin and the time spent there.

### Noise has two competing effects

A purely deterministic description would miss an important feature of the model. Noise is required for barrier crossing when both basins remain stable, but excessive noise also promotes escape from the low-*q* basin. Consequently, the dependence on *D* is non-monotonic.

For *A* = 0.01, increasing *D* from 0.0025 to 0.005, 0.01, and 0.02 increases the low-state probability at drive removal from 0.033 to 0.261, 0.670, and 0.842. A further increase to *D* = 0.04 reduces it to 0.718. After 50 additional time units, the corresponding probabilities are 0.056, 0.358, 0.707, 0.489, and 0.402. Thus, weak noise leaves most trajectories trapped in the initial high-*q* basin, intermediate noise facilitates access to the low-*q* basin, and strong noise destabilizes the resulting low-state population.

This non-monotonicity is one of the most useful predictions of the model because it is difficult to obtain from a purely verbal analogy. In a future empirical mapping, an analogous variable would correspond to a quantified source of trial-to-trial or moment-to-moment neural variability. The present model does not specify which physiological process should play that role. It predicts only that a stochastic system with a metastable low-activity basin can exhibit an intermediate-noise regime in which access is enhanced without complete loss of persistence.

### Occupancy compression and transition entropy are different observables

The original conceptual framework emphasized that stationarity and transition entropy should not be equated with thoughtlessness. The simulations provide a concrete demonstration of this point. As the focusing amplitude becomes strong, the effective number of occupied *q*-bins decreases from approximately *N*_eff_ = 11.95 at *A* = 0.005 to 7.08, 6.33, and 6.31 at *A* = 0.015, 0.020, and 0.030, respectively. In contrast, the transition entropy changes only modestly, from approximately 1.18 at weak drive to 1.16 at the strongest drive.

The behaviour is instructive. A system can concentrate its probability into a restricted region of state space without making every individual transition highly predictable. Conversely, a system can have low transition entropy while continuing to cycle deterministically between states. Therefore, a future neural analysis should not attempt to identify a thoughtless state from one entropy statistic. At minimum, state occupancy, transition structure, persistence, and independent reports of subjective experience should be analyzed jointly.

## Discussion

### A mechanistic interpretation of the computational results

The simulations support a precise version of the proposed driven–relaxation mechanism:

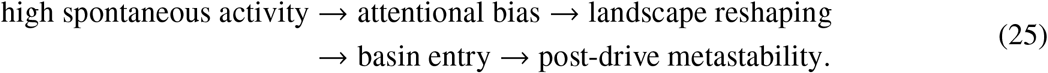

The first arrow is represented by the externally imposed drive *u* (*t*). The second is the coupling between *u* (*t*) and *r* (*t*). The third follows from the tilt term −*rq*. Basin entry occurs through deterministic loss of stability or stochastic barrier crossing, depending on the drive amplitude and noise. Finally, the finite *τ*_*r*_ and the residual landscape structure allow the low-*q* population to persist after drive removal.

The model therefore separates three questions that are often conflated. First, *can a low-activity regime be reached?* The answer is yes for a broad parameter region. Second, *can it persist after the manipulation?* Yes, but only metastably and with a persistence that depends strongly on noise. Third, *does this regime correspond to subjective thoughtlessness?* The model cannot answer this. That last step requires an independent empirical definition of the experience.

### Testable predictions

The computational model converts the conceptual proposal into several predictions that can be evaluated experimentally.

First, if a controlled attentional manipulation can be mapped onto a drive variable, increasing the effective drive should increase the probability of entering a low-spontaneous-thought regime and decrease the first-passage time to that regime. A threshold-like change may occur if the underlying dynamics contain a bifurcation analogous to the spinodal transition.

Second, after the attentional manipulation ends, the candidate state should not necessarily disappear immediately. A transient persistence whose duration depends on the preceding manipulation would support a history-dependent dynamical interpretation. A stimulus-locked response with no post-manipulation persistence would instead favour a simpler driven-response explanation.

Third, the model predicts a trade-off between access and stability as a function of stochastic variability. Too little variability may inhibit transitions into a new basin, whereas excessive variability can destroy its persistence. This prediction is more specific than the generic claim that meditation should reduce or increase neural entropy.

Fourth, occupancy-based and transition-based measures should not necessarily change in the same direction. A candidate low-thought state could show a restricted neural repertoire while retaining substantial transition uncertainty within that repertoire. This provides a direct test of the distinction between state-space compression and transition predictability.

Finally, these predictions should be tested against control conditions. Focused attention, slow breathing, ordinary rest, drowsiness, and other forms of reduced external stimulation may all produce altered neural dynamics. A distinctive signature of the proposed mechanism should therefore be defined relative to these alternatives rather than inferred from a change in one neural statistic alone.

### Limitations and alternative explanations

The model is intentionally minimal and its limitations are therefore substantial. First, *q* is a latent mathematical coordinate, not a validated biomarker of spontaneous thought. The mapping between *q* and subjective experience must be established independently. Second, the double-well landscape is a phenomenological construction. Real neural dynamics are high-dimensional, adaptive, heterogeneous, and coupled across spatial and temporal scales; there is no evidence that the brain possesses a literal potential-energy surface of the form used here.

Third, the present model assumes that focused attention biases the system toward lower spontaneous-thought activity. This sign convention is necessary to define the toy model but is not itself a result. An empirical system could instead show multiple competing effects of attention, arousal, sensory input, and internal regulation. Fourth, the one-dimensional model cannot distinguish different kinds of low-activity states. A low *q* could represent genuine reduction of spontaneous thought, reduced responsiveness, drowsiness, or another dynamical regime unless independent behavioural and subjective measures rule out these alternatives.

Fifth, the numerical results are dimensionless. The values of *A, D, a*, and *τ*_*r*_ should not be interpreted as physiological quantities. The purpose of the simulations is to establish dynamical consequences of the model and to identify qualitative regimes that can motivate experiments. A future data-driven model would need to infer parameters from neural recordings rather than assigning them arbitrarily.

Sixth, entropy estimates depend on coarse-graining. Changing the number of bins or the sampling interval changes both *N*_eff_ and *h*_tr_. This dependence is not a flaw in the information-theoretic definitions, but it means that comparisons across experiments require a predefined and reproducible state-space construction.

Finally, the model does not demonstrate that meditation produces a thoughtless state. It demonstrates only that a driven stochastic system with bistability and a finite response time can produce a low-activity metastable regime after the drive is removed. The scientific value of the model therefore lies in the quantitative predictions it generates, not in treating the analogy as evidence for a particular neural mechanism.

### Scope, falsification, and experimental path

The strongest use of the present model is as a hypothesis-generating and falsifiable framework. A direct experimental test would require time-resolved neural recordings together with an independent identification of periods in which participants report substantially reduced spontaneous thought while remaining awake and responsive. The analysis should compare ordinary rest, focused attention, and the independently identified low-thought condition.

The model would gain support if the low-thought condition showed a reproducible history-dependent transition, increased post-manipulation persistence, and a relationship between the preceding attentional manipulation and the probability of entering the candidate state. A result showing no difference from focused-attention controls would weaken the proposed interpretation. A result showing that the state is entirely stimulus-locked would favour a driven rather than metastable explanation. Conversely, a low-thought state accompanied by increased neural variability rather than reduced variability would not automatically falsify the model; it would falsify only interpretations that equate low thought with low entropy.

The most informative next step is therefore not to search for a universal “neural entropy of thoughtlessness”, but to test the dynamical structure directly: basin entry, transition rates, relaxation times, dwell-time distributions, and their dependence on controlled perturbations. This is where the statistical-mechanical formulation provides a practical advantage over a purely verbal analogy.

## Conclusions

We developed a minimal stochastic statistical-mechanical model for a proposed transition from wandering cognition toward a persistent low-spontaneous-thought regime. The model couples a slowly relaxing attentional-response variable to a tilted double-well landscape and stochastic fluctuations. Numerical simulations show four robust features of the model: a drive-dependent crossover in low-state occupancy, a finite post-drive relaxation period, metastable persistence whose duration depends on noise, and a non-monotonic dependence of state access on stochastic fluctuations.

The model also clarifies a conceptual point that is essential for future empirical work. Stationarity, low occupancy entropy, low transition entropy, and subjective thoughtlessness are distinct quantities. None can be substituted for another. A scientifically useful account must therefore connect model observables to neural measurements and subjective reports without assuming their equivalence.

The present results should not be interpreted as evidence that meditation literally creates a thermodynamic equilibrium or that the brain follows the specific potential used here. They establish a narrower claim: a simple driven stochastic system is mathematically capable of producing the proposed sequence of attentional forcing, basin entry, relaxation, and metastable low-activity residence, and the resulting behaviour is quantitatively sensitive to drive strength and noise. These dependencies generate experimentally testable predictions. If future data fail to show the predicted history dependence or persistence, the model can be rejected or reformulated. If a reproducible dynamical signature is found, the present framework provides a compact starting point for more detailed biophysical and neural models.

## Supporting information

Electronic Supporting Information

## Data Availability

All data generated and analyzed in this study are available with the manuscript. The simulation trajectories, parameter-sweep data, processed observables, and numerical data used to generate the figures are provided as Supporting Information. The Python scripts used to perform the stochastic simulations, calculate the reported observables, and reproduce the figures are also provided to facilitate reproducibility.

## Author information

### Declaration of competing interest

The author declares no conflict of interest.

## Acknowledgments

The author thanks Prime Minister’s Research Fellowship (PMRF) and Department of Chemistry, IIT Kharagpur for funding and support. The author thanks Ramakrishna Mission, Mr. Deepak Khurana, and Ms. Neeranjana Ghosh for introducing him to the field of meditation and mindfulness. The author thanks Prof. Sonjoy Majumdar, Prof. Piyush Mishra, and Mr. Koshis K. Pradhan for useful discussion. This article is dedicated to Prof. Sabyashachi Mishra and Prof. Srabani Taraphdar, who inspired the author to pursue research in quantum chemistry and statistical mechanics.

## Notes

### Competing Interest Statement

The authors have declared no competing interest.

## References

[1] Kalina Christoff, Zachary C. Irving, Kieran C. R. Fox, R. Nathan Spreng, and Jessica R. Andrews-Hanna. Mind-wandering as spontaneous thought: A dynamic framework. Nature Reviews Neuroscience, 17:718–731, 2016.

[2] Susanna Feruglio, Alessio Matiz, Giuseppe Pagnoni, Franco Fabbro, and Cristiano Crescentini. The impact of mindfulness meditation on the wandering mind: A systematic review. Neuroscience and Biobehavioral Reviews, 131:313–330, 2021.

[3] Hadley Rahrig, David R Vago, Matthew A Passarelli, Allison Auten, Nicholas A Lynn, and Kirk Warren Brown. Meta-analytic evidence that mindfulness training alters resting state default mode network connectivity. Scientific Reports, 12(1):12260, 2022.

[4] Thilo Hinterberger, Stephanie Schmidt, Tsutomu Kamei, and Harald Walach. Decreased electrophysiological activity represents the conscious state of emptiness in meditation. Frontiers in psychology, 5:99, 2014.

[5] Mikail Khona and Ila R. Fiete. Attractor and integrator networks in the brain. Nature Reviews Neuroscience, 23:744–766, 2022.

[6] Michael Breakspear. Dynamic models of large-scale brain activity. Nature Neuroscience, 20:340–352, 2017.

[7] Paulina Clara Dagnino, Javier A. Galadí, Estela Càmara, Gustavo Deco, and Anira Escrichs. Inducing a meditative state by artificial perturbations: A mechanistic understanding of brain dynamics underlying meditation. Network Neuroscience, 8(2):517–540, 2024.

[8] Prakash Chandra Kavi, Gorka Zamora-López, Daniel Ari Friedman, and Gustavo Patow. Thoughtseeds: A hierarchical and agentic framework for investigating thought dynamics in meditative states. Entropy, 27(5):459, 2025.

[9] Marieke Van Vugt, Amir Moye, and Swagath Sivakumar. Computational modelling approaches to meditation research: Why should we care? Current Opinion in Psychology, 28:49–53, 2019.

[10] Jonathan Smallwood and Jonathan W. Schooler. The science of mind wandering: Empirically navigating the stream of consciousness. Annual Review of Psychology, 66:487–518, 2015.

[11] Jessica R. Andrews-Hanna, Jonathan Smallwood, and R. Nathan Spreng. The default network and self-generated thought: Component processes, dynamic control, and clinical relevance. Annals of the New York Academy of Sciences, 1316(1):29–52, 2014.

[12] Wendy Hasenkamp, Christine D. Wilson-Mendenhall, Erica Duncan, and Lawrence W. Barsalou. Mind wandering and attention during focused meditation: A fine-grained temporal analysis of fluctuating cognitive states. NeuroImage, 59(1):750–760, 2012.

[13] Judson A. Brewer, Patrick D. Worhunsky, Jeremy R. Gray, Yi-Yuan Tang, Jochen Weber, and Hedy Kober. Meditation experience is associated with differences in default mode network activity and connectivity. Proceedings of the National Academy of Sciences, 108(50):20254–20259, 2011.

[14] Antoine Lutz, Lawrence L. Greischar, Nancy B. Rawlings, Matthieu Ricard, and Richard J. Davidson. Long-term meditators self-induce high-amplitude gamma synchrony during mental practice. Proceedings of the National Academy of Sciences, 101(46):16369–16373, 2004.

[15] Yi-Yuan Tang Britta K. Hölzel, and Michael I. Posner. The neuroscience of mindfulness meditation. Nature Reviews Neuroscience, 16:213–225, 2015.

[16] David R. Vago and David A. Silbersweig. Self-awareness, self-regulation, and self-transcendence (s-art): A framework for understanding the neurobiological mechanisms of mindfulness. Frontiers in Human Neuroscience, 6:296, 2012.

[17] Zoran Josipovic. Neural correlates of nondual awareness in meditation. Annals of the New York Academy of Sciences, 1307:9–18, 2014.

[18] Antoine Lutz, Heleen A. Slagter, John D. Dunne, and Richard J. Davidson. Attention regulation and monitoring in meditation. Trends in Cognitive Sciences, 12(4):163–169, 2008.

[19] Norman A. S. Farb et al. Attending to the present: Mindfulness meditation reveals distinct neural modes of self-reference. Social Cognitive and Affective Neuroscience, 2(4):313–322, 2007.

[20] Michael D. Mrazek, Michael S. Franklin, Dawa Tarchin Phillips, Benjamin Baird, and Jonathan W. Schooler. Mindfulness training improves working memory capacity and gre performance while reducing mind wandering. Psychological Science, 24(5):776–781, 2013.

[21] Stanislas Dehaene and Jean-Pierre Changeux. Experimental and theoretical approaches to conscious processing. Neuron, 70(2):200–227, 2011.

[22] Giulio Tononi, Melanie Boly, Marcello Massimini, and Christof Koch. Integrated information theory: From consciousness to its physical substrate. Nature Reviews Neuroscience, 17:450–461, 2016.

[23] Karl Friston. The free-energy principle: A unified brain theory? Nature Reviews Neuroscience, 11: 127–138, 2010.

[24] Lars Sandved-Smith. A computational model of minimal phenomenal experience (mpe). 2025.

[25] Nike Walter and Thilo Hinterberger. Determining states of consciousness in the electroencephalogram based on spectral, complexity, and criticality features. Neuroscience of Consciousness, 2022(1):niac008, 2022.

[26] R. Martínez Vivot et al. Meditation increases the entropy of brain oscillatory activity. Neuroscience, 431:40–51, 2020.

[27] Daniel A Atad, Pedro AM Mediano, Fernando E Rosas, and Aviva Berkovich-Ohana. Meditation and complexity: a review and synthesis of evidence. Neuroscience of Consciousness, 2025(1):niaf013, 2025.

[28] Bill Lubanovic. Introducing Python: modern computing in simple packages. “ O’Reilly Media, Inc.”, 2019.

