## Supplementary material for "From Wandering Thoughts to Mental Stillness: A Minimal Computational Statistical-Mechanical Model of Driven Relaxation": Electronic Supporting Information

### 1 Mathematical Background

The model combines deterministic dynamical systems, stochastic differential equations, and information-theoretic measures. The mathematical framework used in the simulations is summarized below.

#### 1.1 Dynamical System and Double-Well Potential

The state of the system is described by a dimensionless coordinate $q(t)$, which represents a coarse-grained measure of spontaneous-thought activity. Its deterministic dynamics are governed by the gradient of a potential $U(q;r)$,

$$\frac{dq}{dt} = -\frac{\partial U(q;r)}{\partial q}. \tag{1}$$

The potential used in the present work is

$$U(q;r) = \frac{a}{4}(q-q_L)^2(q-q_H)^2 - rq, \tag{2}$$

where $q_L$ and $q_H$ define the low- and high-$q$ regimes, $a$ controls the barrier between them, and $r$ introduces a time-dependent tilt. The stationary points satisfy

$$\frac{\partial U}{\partial q} = 0, \tag{3}$$

with minima corresponding to locally stable states and the intervening maximum representing the barrier.

For Eq. 2,

$$\frac{\partial U}{\partial q} = a(q-q_L)(q-q_H)(2q-q_L-q_H) - r. \tag{4}$$

#### 1.2 Stochastic Dynamics

Random fluctuations are incorporated through a Langevin equation,

$$\frac{dq}{dt} = -\frac{\partial U(q;r)}{\partial q} + \sqrt{2D}\,\xi(t), \tag{5}$$

where $D$ is the noise intensity and $\xi(t)$ is Gaussian white noise satisfying

$$\langle \xi(t) \rangle = 0, \qquad \langle \xi(t)\xi(t') \rangle = \delta(t - t'). \tag{6}$$

Thus, the first term determines the deterministic drift on the potential landscape, whereas the second term produces stochastic barrier crossing.

### 1.3 History-Dependent Drive and Relaxation

The tilt variable $r(t)$ evolves according to

$$\frac{dr}{dt} = -\frac{r}{\tau_r} + \alpha u(t), \tag{7}$$

where $\tau_r$ is the relaxation time, $\alpha$ is the coupling between the external input and the tilt, and $u(t)$ represents the focusing protocol. The finite-duration drive is defined as

$$u(t) = \begin{cases} -A, & 0 \le t < t_f, \\ 0, & t \ge t_f, \end{cases} \tag{8}$$

where $A$ is the drive amplitude and $t_f$ is the drive-removal time.

After drive removal, $u(t) = 0$, giving

$$r(t) = r(t_f) \exp\left[-\frac{t - t_f}{\tau_r}\right]. \tag{9}$$

Thus, the slow variable retains a transient dependence on the preceding focusing period and gradually relaxes toward zero.

### 1.4 Spinodal Condition

Increasing the magnitude of $r$ progressively destabilizes one of the potential wells. The deterministic spinodal point is obtained when a minimum and the neighboring maximum merge, satisfying

$$\frac{\partial U}{\partial q} = 0, \qquad \frac{\partial^2 U}{\partial q^2} = 0. \tag{10}$$

For

$$d = \frac{q_H - q_L}{2}, \tag{11}$$

the corresponding critical tilt is

$$|r_{\mathrm{sp}}| = \frac{2ad^3}{3\sqrt{3}}. \tag{12}$$

Since the steady-state magnitude of $r$ under a constant drive is $\alpha A \tau_r$, the corresponding drive scale is

$$A_{\mathrm{sp}} = \frac{|r_{\mathrm{sp}}|}{\alpha \tau_r}. \tag{13}$$

For the parameterization used here, $A_{\mathrm{sp}} \simeq 0.0148$. This value is a model-dependent dimensionless threshold and has no direct physiological interpretation.

### 1.5 State Occupancy and First-Passage Quantities

The low-$q$ state is defined using a threshold $q_{\text{th}}$, such that

$$q < q_{\text{th}}. \tag{14}$$

For an ensemble of $N$ independent trajectories, the probability of occupying the low-$q$ regime at time $t$ is

$$P_L(t) = \frac{N_L(t)}{N}, \tag{15}$$

where $N_L(t)$ is the number of trajectories satisfying the threshold condition.

The first-passage time is the time required for a trajectory to reach the low-$q$ threshold for the first time,

$$T_{\text{FP}} = t_{\text{FP}} - t_0, \tag{16}$$

and its ensemble mean is

$$\langle T_{\text{FP}} \rangle = \frac{1}{N} \sum_{i=1}^{N} T_{\text{FP}}^{(i)}. \tag{17}$$

Similarly, if a trajectory enters and leaves the low-$q$ region at $t_{\text{in}}$ and $t_{\text{out}}$, respectively, its dwell time is

$$T_{\text{dwell}} = t_{\text{out}} - t_{\text{in}}. \tag{18}$$

### 1.6 Numerical Integration

The stochastic differential equation is integrated using the Euler–Maruyama scheme. For a time step $\Delta t$,

$$q_{n+1} = q_n - \left.\frac{\partial U}{\partial q}\right|_{q_n, r_n} \Delta t + \sqrt{2D\Delta t}\, \eta_n, \tag{19}$$

where

$$\eta_n \sim \mathcal{N}(0, 1). \tag{20}$$

The relaxing variable is propagated as

$$r_{n+1} = r_n + \left( -\frac{r_n}{\tau_r} + \alpha u_n \right) \Delta t. \tag{21}$$

Independent stochastic trajectories are generated using independent realizations of $\eta_n$.

### 1.7 Probability Distributions and Entropy

For information-theoretic analysis, the continuous $q$-coordinate is discretized into $M$ bins. If $p_i$ denotes the probability of occupying bin $i$,

$$\sum_{i=1}^{M} p_i = 1. \tag{22}$$

The occupancy entropy is calculated using the Shannon entropy,

$$H_{\text{occ}} = -\sum_{i=1}^{M} p_i \ln p_i. \tag{23}$$

The corresponding effective number of occupied states is

$$N_{\text{eff}} = \exp(H_{\text{occ}}). \tag{24}$$

A lower $N_{\text{eff}}$ therefore indicates that the probability distribution is concentrated over fewer $q$-states.

### 1.8 Transition Probabilities and Transition Entropy

The discretized trajectories also define a transition matrix $\mathbf{P}$, where

$$P_{ij} = P(X_{t+\Delta t} = j \mid X_t = i), \tag{25}$$

and

$$\sum_{j=1}^{M} P_{ij} = 1. \tag{26}$$

For a stationary state distribution $\boldsymbol{\pi} = (\pi_1, \dots, \pi_M)$,

$$\boldsymbol{\pi}\mathbf{P} = \boldsymbol{\pi}. \tag{27}$$

The transition entropy is then

$$H_{\text{tr}} = -\sum_{i=1}^{M} \pi_i \sum_{j=1}^{M} P_{ij} \ln P_{ij}. \tag{28}$$

Occupancy entropy and transition entropy quantify different aspects of the dynamics: $H_{\text{occ}}$ measures the diversity of occupied states, whereas $H_{\text{tr}}$ measures the uncertainty of subsequent transitions. Consequently,

$$H_{\text{occ}} \neq H_{\text{tr}}, \tag{29}$$

and neither quantity alone is sufficient to characterize dynamical "stillness."

### 1.9 Statistical Averaging

For an observable $X$ obtained from $N$ independent trajectories, the ensemble mean is

$$\langle X \rangle = \frac{1}{N} \sum_{i=1}^{N} X_i. \tag{30}$$

The sample standard deviation and standard error of the mean are

$$s = \sqrt{\frac{1}{N-1} \sum_{i=1}^{N} (X_i - \langle X \rangle)^2}, \tag{31}$$

and

$$\text{SEM} = \frac{s}{\sqrt{N}}, \tag{32}$$

respectively.

All variables in the present model are dimensionless; consequently, the numerical values of $q$, $r$, $A$, $D$, and time are model units and are not directly assigned physiological units.
